# Automated Processing and Statistical Analysis of NMR spectra obtained from *Arabidopsis thaliana* Extracts

**DOI:** 10.1101/766295

**Authors:** Valtteri Mäkelä, Lauri Vaahtera, Jussi Helminen, Harri Koskela, Mikael Brosché, Ilkka Kilpeläinen, Sami Heikkinen

**Author notes:** Corresponding author Email address (Valtteri Mäkelä).

## Abstract

To cope with vast amounts of data produced by metabolomic and mixture analysis using NMR, the employed analysis techniques and tools are very important. In this paper, we demonstrate quantitative ^1^H and 2D-JRES NMR analysis of *Arabidopsis thaliana* extracts utilizing our ImatraNMR and SimpeleNMR software to automate the processing the spectra, extracting data, and perform statistical analysis of the results. Metabolomes of four different strains of *Arabidopsis thaliana* are analyzed under control conditions and during acute ozone exposure. Key differences are identified between accessions Col-0 and Cvi-0 (epithionitriles, iberin nitrile) and ozone damage associated metabolite(s) can be identified. Sample stability is also addressed.

## 1. Introduction

Quantitative techniques and mixture analysis with NMR are becoming increasingly common, while high-field NMR instruments, cryoprobes and autosamplers are available to more researchers. Metabolite analysis, reaction monitoring/following or other mixture analysis can produce vast amounts of complex spectra, which are difficult to analyze with traditional NMR processing software, as they are designed from the viewpoint of small-molecule analysis. To accommodate these needs, features aimed for mixture analysis and automation have been incorporated into commercial processing software (Agilent/Varian CRAFT [1], Mnova GSD and qNMR [2], Bruker Biospin AMIX and IconNMR). The rising popularity of utilizing NMR for metabolomics has also spawned many commercial and free tools suited for different stages and aspects of metabolite analysis: NMR Suite (Chenomx Inc. [3, 4]), rNMR [5], BATMAN [6, 7], Pathomx [8] and others [9, 10], with publicly available databases [11, 12] and even web-applications available [13, 14].

Even with these developments, the workflow for processing and analyzing hundreds of spectra can be a complex, and purpose-built tools might be suitable only for certain type of analysis, be vendor specific or expensive. More general purpose raw processing tools such as NMRPipe [15] and NMRGlue [16] can be used in many tasks, but require scripting or programming knowledge from the user. To address some aforementioned limitations, we created simple, free, and multi-vendor processing/analysis software ImatraNMR and SimpeleNMR. These tools are more generic and suitable for automation than traditional processing programs, but are still somewhat simpler to use than raw processing tools and provide data-analysis capabilities. While originally developed for automated hydrocarbon analysis[17], in this paper we use updated versions of these tools and demonstrate metabolite analysis of *Arabidopsis thaliana* extracts, and discuss various difficulties and techniques associated with this type of analysis.

### 1.1. Sensitivity and overlap

Metabolite analysis and mixture analysis in general using NMR has two main obstacles: sensitivity and signal overlap. The most sensitive basic technique, 1D ^1^H NMR, is simple to acquire and quite easily quantitative, but suffers from low chemical shift dispersion and signal splitting due J-coupling. Multidimensional NMR techniques such as 2D HSQC provide significant boost in resolving power, but come with the cost of reduced sensitivity and/or increased acquisition time. Multipulse experiments are also more complex and depend on a large number of phenomena, including usually magnetization transfer, rendering most experiments non-quantitative without careful preparation and/or redesign of the experiment.

If sample concentration is high enough (such as in hydrocarbon analysis), even quantitative 1D ^13^C NMR can be used, as it is simple to acquire and provides much greater chemical shift dispersion [17]. While usually not accessible, it can be viable even in metabolomic context if hyperpolarization can be utilized [18]. Essentially limitations in resolving power can be generally transformed to limitations in sensitivity, providing that the spectrometer at hand can implement modern NMR techniques.

The resolving power of ^1^H spectra can also be improved by eliminating homonuclear ^1^H -^1^H couplings, rendering all signals to singlets. This is not trivially done, but several modern techniques such as those based on Zangger-Sterk element are capable of creating homonuclearly decoupled or *pure shift* ^1^H spectrum [19, 20], details of which can be found in recent review by Zangger [21]. Unfortunately most techniques suffer from prohibitive sensitivity penalty of retaining only few percents of the signal, making them difficult to utilize in metabolic analysis. The classic J-resolved spectra (2D-JRES)[22] processed appropriately can also yield essentially decoupled spectra with less signal loss, while quantitativity is sacrificed by the use of sine-bell type apodization along absolute value processing to avoid dispersive line shape. There are techniques which can be used to circumvent this, such as advanced time-domain processing [23, 24, 25, 26, 27] and the use of Zangger-Sterk element [28], but they greatly increase the complexity of the processing, or again suffer from significant sensitivity penalty.

### 1.2. Quantitative NMR

Traditionally, quantitative NMR means that all signal intensities are directly proportional to concentration, and the absolute concentration can be easily derived from known concentration of internal/external standard. These quantitative conditions are easily achievable on basic single pulse 1D spectra, and they are possible in some specialized versions of more complex experiments such as the quantitative versions of INEPT [29, 30], DEPT/POMMIE [31, 32] and HSQC [33, 34, 35]. In other experiments, non-uniform factors affecting signal intensity makes straightforward comparison impossible and calibration curves and authentic standards are required, which can be difficult or infeasible.

This drawback doesn’t render the experiments unsuitable for quantitative analysis as such, as the signal intensity can usually assumed to be linearly proportional to concentration when measuring all samples under identical conditions and field strength. So while all signals can’t be directly compared to each other, signals originating from same compound can still be compared across numerous samples. This enables studying the relative concentration differences, which in many cases is enough, especially in metabolite analysis where the absolute concentration might anyway depend strongly on sample preparation or other factors.

### 1.3. NMR processing and data extraction

To produce meaningful data, the acquired FIDs must be processed and data extracted from the resulting spectra, preferably via automatic, consistent and robust process when large numbers of spectra are involved. Luckily, almost all of the processing parameters can be usually predetermined or devised easily from the experimental setup for a set of similar spectra, at least after looking at few representative samples. Notable exception is phase correction, which still is required to be performed manually in some cases. Most phase problems are related to delayed sampling, transient response of analog filters and errors in the first few data points, which can lead to significant baseline errors after phase correction [36, 37, 38]. Recent spectrometers with very high oversampling rates and DSP filtering (Agilent VNMRS/DD2 consoles) [39, 40, 41] and/or backward prediction of the first few data points (TopSpin digitizer mode “BASEOPT” [42, 43]) have diminished these problems and generally produce spectra with good baseline and low 1st order phase correction. Still, some error sources such as phase response of RF pulses can’t be rectified with these measures, and a totally automatic and robust method suitable for every case seems to be difficult to devise. Subsequently, new methods have been developed even quite recently [44, 45]. Phasing can be also avoided with absolute value processing which renders phase meaningless, but this has severe drawbacks as discussed below.

After processing the signals must be quantified for analysis, which is easy for few well separated signals using numerical integration, but rapidly tedious when dealing with hundreds of spectra without proper automation. Overlapping signals can be separated with line shape fitting quite accurately, but automation is still limited and the technique is usually only employed in more targeted analysis, when the signals resulting from compounds of interest are known and can be fitted simultaneously [46, 4, 47, 6, 7]. Still, impressive automation can be achieved with tailored analysis tools for specific class of samples: for example fully automated analysis of blood plasma has been demonstrated utilizing advanced statistical methods such as Bayesian models, Markov Chain Monte Carlo (MCMC) and Sequential Monte Carlo (SMC) [48, 49, 13].

Other techniques like Agilent CRAFT [1] based on Bayesian fitting of signals directly to the FID data or Mnova GSD [2] can in principle decompose the spectrum into frequency-intensity pairs in full automation, but do not take into account or combine signals resulting from the same compound. This kind of data requires further refinement or filtering of the data in non-targeted analysis, and the real-world use of these techniques applied for complex mixtures seems to be limited.

If no particular compounds are targeted, the spectrum area is usually divided in equal integration areas, “bins”, which are integrated and treated as variables describing the sample and analyzed with statistical software. In addition to traditional equidistant binning, several more sophisticated schemes have been developed [50, 51, 52], with the aim of more meaningful placement of the bin borders. In ImatraNMR, simple scheme based on classifying found signals can be used (“histogram binning”).

### 1.4. Metabolite analysis and JRES

While the 2D J-resolved spectroscopy (2D-JRES) as a technique is quite old [22], it is suitable for metabolomic analysis in many ways, as discussed in review article of Ludwig et al.[53]. JRES spectrum is processed in absolute value mode, so phasing is not required at all, making the automation of processing the spectrum straightforward and robust, even in the presence of broad signals or water suppression residual. The broadening caused by dispersive line shape is minimized by sine bell apodization, while shearing, symmetrization and projection post-processing neutralize the effect of ^1^H -^1^H couplings producing essentially “decoupled” spectra. The drawback of the sine window function is the strong weighting of signal intensity by T_2_ relaxation rate, which compromises absolute quantitativity. However this is not essential when comparing relative concentrations, and JRES has been established as routine tool in metabolite analysis. If needed, a calibration curve can be used producing accuracy and precision can be comparable to ^1^H NMR spectra [54].

The shear/rotate and symmetrization operations can produce some artifacts, however they seem to be a minor problem in practice: Parsons et al. have studied the quantification errors with real and simulated JRES spectra, and found that the post-processing causes less than 2% of errors in quantification [55]. Signals in close proximity can cause significant errors up to about 33%, but this error falls rapidly to under few percent with distance. Furthermore, in real-world application to metabolite analysis, the errors were comparable to those obtained by line shape fitting of ^1^H spectra [55].

Reducing the proton signals to singlets also simplifies analysis and integration: the spectrum will have smaller number signals, and if binning is used, the narrower signal footprint reduces the probability of a signal being spread into several bins. This can be expected work well in combination with non-equidistant binning schemes such as the “seek histogram” feature in ImatraNMR.

### 1.5. SimpeleNMR and ImatraNMR analysis software

SimpeleNMR can perform the basic processing of the FID using discrete Fourier transform (DFT) to produce spectrum, along with the usual pre- and post-processing steps (apodization, baseline correction). ImatraNMR on the other hand is aimed for batch integration of NMR spectra, with some preprocessing functionality, and can read spectra in multiple formats as well. Both programs are freely available at http://vltr.fi/imatranmr/, and can be freely used for research and commercial purposes.

Recently, some new features have been added for both programs, making the them more suitable for metabolite analysis. In SimpeleNMR, 2D-JRES processing has been implemented, with limited support for other magnitude mode 2D spectra. The usual post-processing techniques of 45°rotation/shear and symmetrization in F2 dimension are implemented as well, allowing fast batch processing into projected 1D spectra. Furthermore, more window functions, improvements in plotting/viewing, supported output formats and basic multivariate (PCA, PLS-DA) analysis based on Scikit-learn [56] have been included. In ImatraNMR, probabilistic quotient normalization by Dieterle et al. has been implemented [57], which can be used for normalization in complex biological samples, where the concentration of the samples is hard to determine. Most of these features are demonstrated in the analysis below.

## 2. Materials and methods

### 2.1. Plant lines

The analysis used 4 plant lines: Natural accessions Col-0 (ozone tolerant) and Cvi-0 (ozone sensitive), and near-isogenic lines Col-s and Cvi-t, where the ozone phenotype of the other natural accession was introgressed into the other. Additionally, 6 recombinant inbred lines (RIL) were analyzed. Of these, four different sample sets were measured. The features are described in Table 1, while the sample sets are described in Table 2.

**Table 1:** The used plant lines.

| Plant line | Description |
| --- | --- |
| Col-0 | Ozone tolerant natural accession |
| Cvi-0 | Ozone sensitive natural accession |
| Col-s | Cvi-0's ozone sensitivity in Col-0 background |
| Cvi-t | Col-0's ozone sensitivity in Cvi-0 background |

**Table 2:** The analyzed sample sets.

| Series | Description | Plant lines |
| --- | --- | --- |
| 1A | First growth period | Col-0, Cvi-0, Col-s, Cvi-t |
| 1B | Second growth | Col-0, Cvi-0, Col-s, Cvi-t |
| 2 | Ozone stress | Col-0, Cvi-0, Col-s |
| 3 | Recombinant inbred lines | Col-0, Cvi-0 + 6 RILs |

Plants were grown on 1:1 peat:vermiculite mixture in controlled environment chambers (Weiss Bio 1300) with 12-h/12-h (day/night) cycle, temperature 22/19 °C, and relative humidity of 70/90 % until the age of three weeks. Plants were collected at 2 pm, after 7 hours of light. Ozone treatment was performed using two similar chambers, other containing 350 ppb ozone. Plants were transferred from the fresh air chamber to the ozone chamber 120 minutes, 60 minutes, 40 minutes, or 20 minutes before 2 pm. Then all samples were collected at 2 pm and flash frozen in liquid nitrogen.

### 2.2. Preparation of NMR samples

The plants were harvested and placed in liquid nitrogen, and ground into fine powder in a mortar cooled with liquid nitrogen. To remove water, the material was lyophilized for 40 hours. For each sample, 25 mg of dried material were weighted and extracted with 1.0 ml of NMR solvent, consisting of 2:8 (vol.) mixture of MeOH-d4 and D_2_O and containing 0.05% TSP-d4 as internal standard. The sample was mixed by vortexing for 30 seconds, heated to +50 ° C for 10 minutes, vortexed 2*10 s during the heating and 30 s after the heating. Then sample was centrifuged at 30000 g for 5 minutes at +4 °C and the supernatant was added to standard 5 mm NMR tubes. The samples were kept on ice until measurement within a few hours. The preparation protocol was adopted from earlier work [58, 59].

### 2.3. NMR spectroscopy

All spectra were acquired at 27 °C on a Varian ^UNITY^INOVA 600 MHz NMR spectrometer using 5 mm triple resonance (^1^H, ^13^C, ^15^N) pulsed field gradient probehead. From every sample, a quantitative ^1^H spectrum was measured using 45° excitation pulse, 13.0 s relaxation delay and 2.0 s of acquisition time, based on maximum observed T_1_ values of about 5 s. Initially, 512 transients were collected (sample series 1A), but for all subsequent samples 256 transients were used as the signal-to-noise ratio seemed sufficient, yielding total experiment time of 1 hour 4 minutes. Pulse length for 90° pulse was calibrated to be ∼ 7.45 us, spectrum window was 7165 Hz (11.94 ppm), while the transmitter was centered at 4.18 ppm.

From sample set 2 (oxidative stress), 2D J-resolved (JRES) spectra were obtained using the same spectrum window and transmitter position, but with 5 s relaxation delay and 1 s acquisition time. In indirect dimension, 64 increments were collected, and for each increment, 8 transients were used, yielding total experiment time of 56 minutes. Additionally, HSQC, HMBC, TOCSY and COSY experiments were measured from select samples to aid in metabolite assignment.

To investigate sample stability, two samples were measured for extended periods of 44 and 160 hours in 27 °C, acquiring ^1^H spectrum for every ∼ 15 minutes. The same parameters were used as in regular quantitative ^1^H experiments, but only 64 transients were collected to achieve the shorter acquisition time.

### 2.4. NMR processing (SimpeleNMR)

All spectra were processed using SimpeleNMR. For regular ^1^H spectra, FIDs were zerofilled doubling the number of data points, and apodized by exponential decay function producing linear broadening of 0.5 Hz. After Fourier transform, automatic phasing (autophase mode 4) was applied, and 1st order baseline correction (drift correction) performed. For JRES, zerofill was performed identically, but a sine bell apodization was used matched to the length of the FID. As typical with JRES, 45 degrees rotation/shear and symmetrization in indirect dimension was performed, and both 2D and 1D versions (via skyline projection of the spectra) were produced. The exact configuration files used with SimpeleNMR can be found in supporting information.

### 2.5. Data extraction (ImatraNMR)

Further processing and extraction of integrals were carried out in ImatraNMR. The processing “recipes” are determined by ImatraNMR script files, which can be found in the supporting information to provide more details. First all spectra were referenced and normalized to the TSP-d4 signal present, which was aligned to 0.0 ppm. Methanol and water residual signals were removed from the spectrum using the nuke function, and signal seek was performed in the region of 0.5-9.0 ppm. From the results, “signal histogram” was generated combining similar signals, and the resulting integral areas integrated (histogram binning). In this method, close signals are classified into same bins and the integral area is determined from the included signals. This avoids including areas lacking detected signals, and produces integration areas which more closely match the present signals compared to equidistant binning. It is also usable in 2D spectra, for which equidistant binning becomes impractical, at least without filtering bins actually containing meaningful signals. For comparison, equidistant binning with 700 regions in the interval of 0.5 - 9.0 ppm was also performed.

While the TSP-d4 acted as internal standard with constant concentration compared to plant material, probabilistic quotient normalization [57] implemented in ImatraNMR was used to generate alternative data sets which does not rely on TSP-d4 concentration.

### 2.6. Statistical analysis

The bin integral files produced by ImatraNMR were loaded easily into the R statistical package [60], in which several PCAs (Principal Component Analysis) were performed for the different sample sets [61]. PCA is a statistical method which decomposes the data into *principal components* of decreasing importance, each one describing part of the variance between samples. The meaning of the components can be interpreted from the the component loading coefficients, which correspond to integral areas and subsequently chemical shifts of distinct metabolites. The highest loads and their bin numbers were extracted from the results and plotted with SimpeleNMR *simp_view.py* to visualize the related signals.

At later stage basic PCA and the related PLS/PLS-DA analysis tools were incorporated to SimpeleNMR using Scikit-learn [56, 62] (*simp_mvar.py*), and the analysis was duplicated. In addition, PLS analyses were conducted for the ozone treatment samples (Set 2) using the exposure time as a response variable.

### 2.7. Metabolite assignment

Most of the metabolites were assigned with BMRB and SDBS databases [11, 63], previous literature concerning similar samples[64, 65, 66] and HSQC, HMBC and COSY/TOCSY obtained from select samples. Epithionitriles and iberin nitrile were not found in the databases, and were simulated by ACD/NMR Predictors 2012 software to obtain reference spectra. A spectrum of C5 epithionitrile was obtained from Koichiro Shimomura [66], but it was also decided to synthesize it as a reference compound, as it was a central differentiator (Figure 1). The synthesis presented in supplementary material. During the assignment process, SimpeleNMR viewer *simp_view.py* was used to overlay spectra, and correlation group tool (*simp_corrg.py*) was used to search for bins whose intensity is collinear with a bin of interest. This is not direct evidence, but can give hints which signals are related even without correlations in 2D spectra, similarly to STOCSY[67].

**Figure 1:**
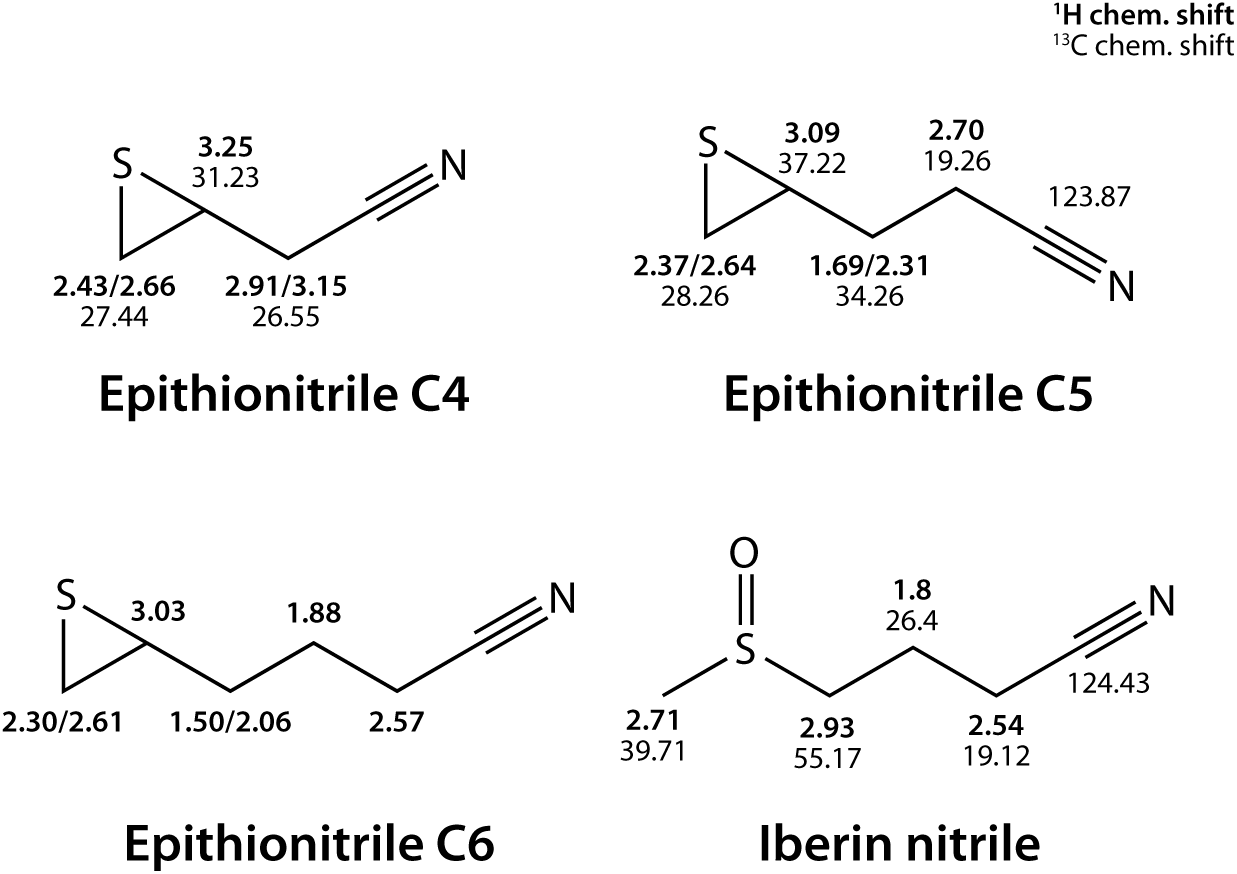
The structure and chemical shift assignments for epithionitriles and iberin nitrile. The iberin nitrile assignments are from a plant extract sample.

## 3. Results and discussion

### 3.1. Separation of plant lines (set 1A and 1B)

The initial sample set included four biological replicates of four different plant lines, however the repetitions were grown in two different time periods, yielding sample sets 1A and 1B. This is reflected in the PCA results (Figure 2), where the first principal component (PC1) separates the sample sets, while the second (PC2) separates the plant lines. The difference between sample sets seems to be related to citric acid cycle, which apparently is more active in 1A. To emphasize the metabolic differences between plant lines, PCA can be performed separately for only 1A or 1B. This approach can be seen in the lower PCA plot, in which the PC1 separates the plant lines already, while PC2 separates the Col-0 and Col-s species further.

**Figure 2:**
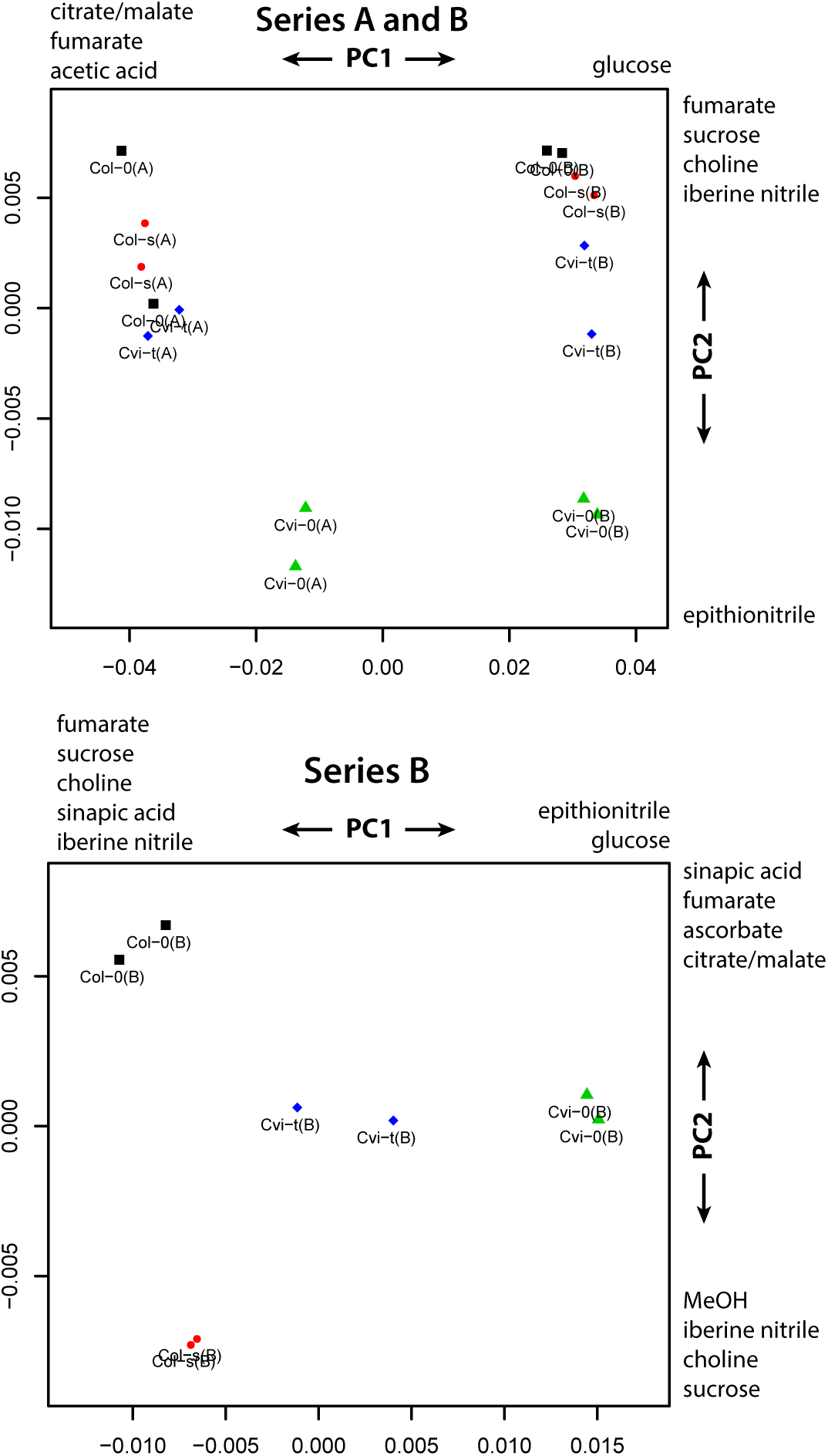
The basic PCA analysis of the four different plant lines, including sample sets 1A and 1B (top), and only 1B (bottom). The growing period can be separated easily (top) while plant lines are separated nicely when analyzing each set individually. All data presented is regular 1H spectra with ImatraNMR histogram binning and PQ normalization.

The main difference between Col-0 and Cvi-0 samples is the C4/C5 epithionitrile and iberin nitrile concentration (Figure 1), which appears to be the characteristic and easily detected distinction, even just by comparing the spectra visually (Figure 3). Differences between Col-0 and Col-s seem to be related to sinapic acid, methanol and citric acid cycle. The C5 epithionitrile and iberin nitrile seem to be result of alternate pathways, and mostly one size was found to be present (Figure 3).

**Figure 3:**
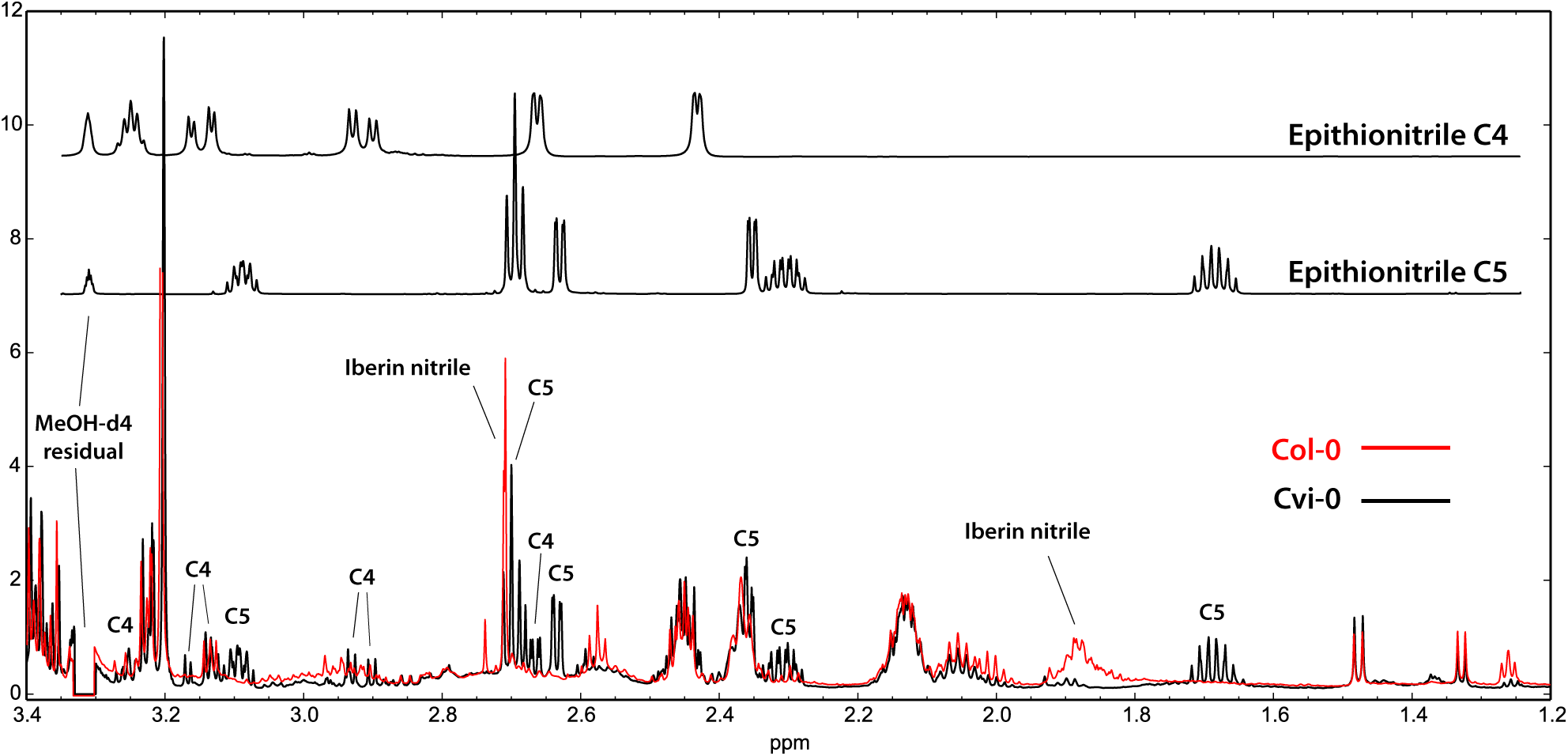
The dissimilarity of Col-0 and Cvi-0 samples can be seen easily by comparison of the ^1^H spectra, as epithionitriles and iberin nitrile have several characteristic signals. Especially the C5 epithionitrile signal at ∼1.7 ppm is easy to observe.

### 3.2. Ozone treatment (set 2)

The Cvi-0, Col-0 and Col-s strains exposed to ozone for 0, 20, 40, 60 and 120 minutes were analysed with both regular ^1^H NMR spectra and JRES spectra, as more subtle differences were expected. The resulting PCA analysis obtained from JRES is presented in Figure 4. Unsurprisingly, the first component separates Col-0 and Cvi-0 strains with similar metabolites as in the first analysis, but PC2 is clearly related to ozone exposure. Short ozone exposure seems to be related with (small) positive score and long exposure with negative score on PC2. The negative PC2 loadings are associated with GABA (Gamma-Aminobutyric acid) and various amino acids, and the GABA concentration indeed rises sharply in 120 min samples in both strains showing visible cell death in that time point (Cvi-0 and Col-s), but not in ozone tolerant Col-0 (Figure 5). This is clearly reflected in the PCA results, where 120 min samples of Col-s and Cvi-0 have much higher negative score on PC2. The rise in the GABA levels coinciding with ozone-induced cell death was an intriguing finding, since Arabidopsis gene GAD4 (AT2G02010), one of the five glutamate decarboxylases catalysing the final step in GABA biosynthesis, has been reported to be one of the most highly ozone-induced transcripts [68]. This implies that GABA might play a significant role during ozone-induced programmed cell death.

**Figure 4:**
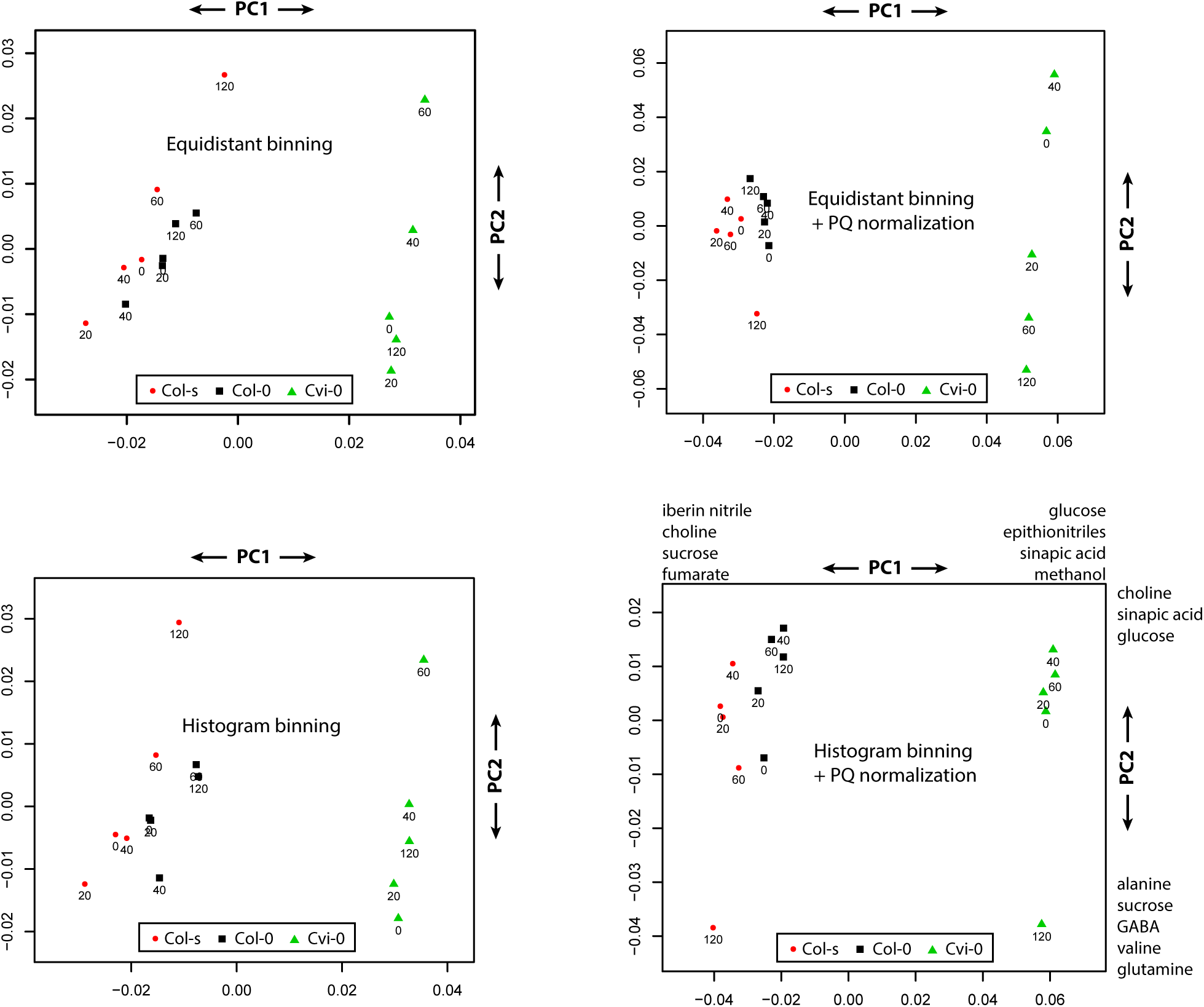
The PCA created from 1D JRES of ozone treated samples, with the exposure time noted in label. Ozone tolerant Col-0 species (black square) can be differentiated easily. The “histogram binning” combined with PQ normalization clearly produces best separation.

**Figure 5:**
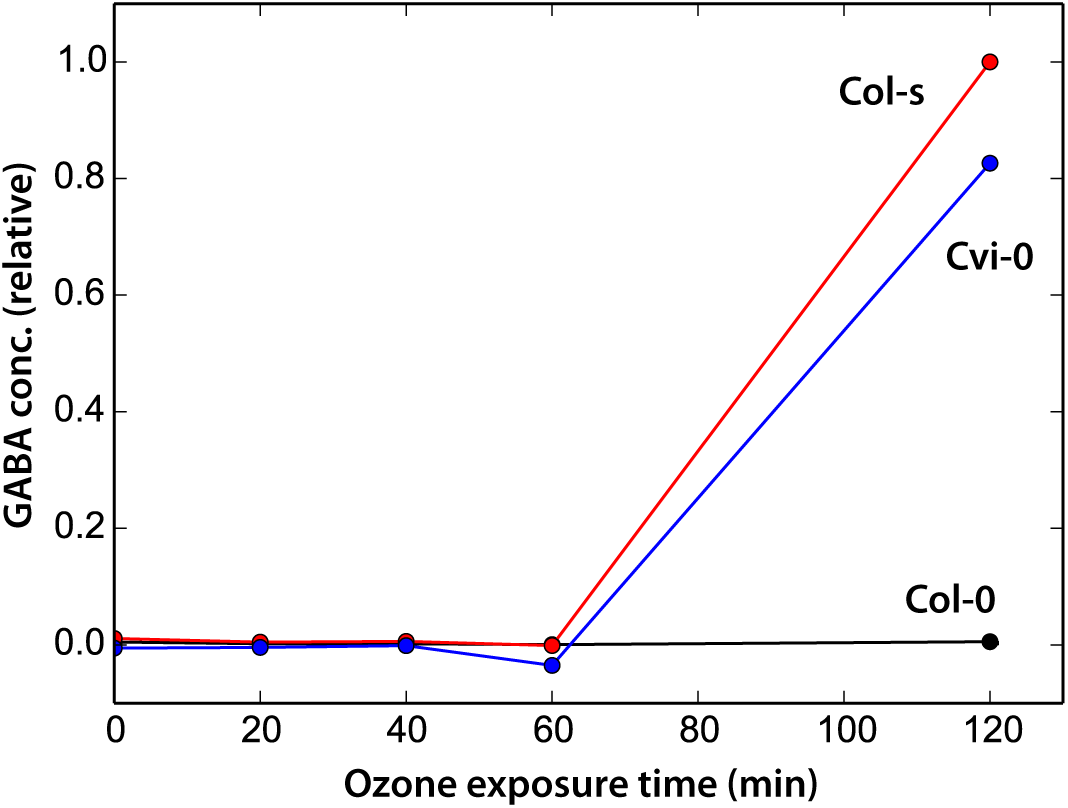
The relative concentration of GABA during ozone exposure (3.01 ppm CH_2_, JRES). The concentration is effectively below detection limit until 60 min, after which it is observed only in the ozone sensitive lines Cvi-0 and Col-s.

In this analysis, the impact of using PQ normalization (as opposed to TSP-d4 internal standard) and ImatraNMR histogram binning (as opposed to equidistant binning) can be demonstrated. As seen in Figure 4, using only histogram binning or equidistant binning with PQ normalization produce inferior results when compared with combined use of both histogram binning and PQ normalization. Histogram binning alone produces similar results to equidistant binning, but works much better in conjunction with PQ normalization, showing clear separation of 120 min samples. Similar results could be achieved with PLS analysis of the same JRES data, but it also revealed the somewhat rising MeOH concentration corresponding to ozone exposure time (non-deuterated MeOH can be separated from the solvent residue due to slight difference in chemical shift). PCA analysis of regular ^1^H derived data also produced similar results, while the improvement provided by histogram binning and PQ normalization was milder. Additional plots can be found in supplementary material.

### 3.3. Sample stablity

Obtaining repetitive quantitative ^1^H spectra for extensive period of time produces a data set which ImatraNMR and SimpeleNMR are well suited: a large number of similar and complex spectra, which must be processed identically and examined.

The two samples were analyzed with PCA similarly as the sample sets 1A/1B and 2, producing a quite interesting plot (Figure 6). The spectra is separated nicely by time by PC1, while PC2 contains much smaller scores. From the the corresponding metabolites of PC1, it can be clearly seen that sucrose and ascorbic acid are decomposing, which can be confirmed easily by looking the spectra and integrals directly (Figure 6). The increase of glucose and fructose are also in line with this observation, suggesting slow hydrolysis of sucrose, and the reaction seems also approximately follow first order kinetics, with a half life of 25.5 hours. Other metabolites seem to be stable, and even the unstable metabolites seem to have half-life of more than >20 h, making the decomposition during the acquisition time relatively small.

**Figure 6:**
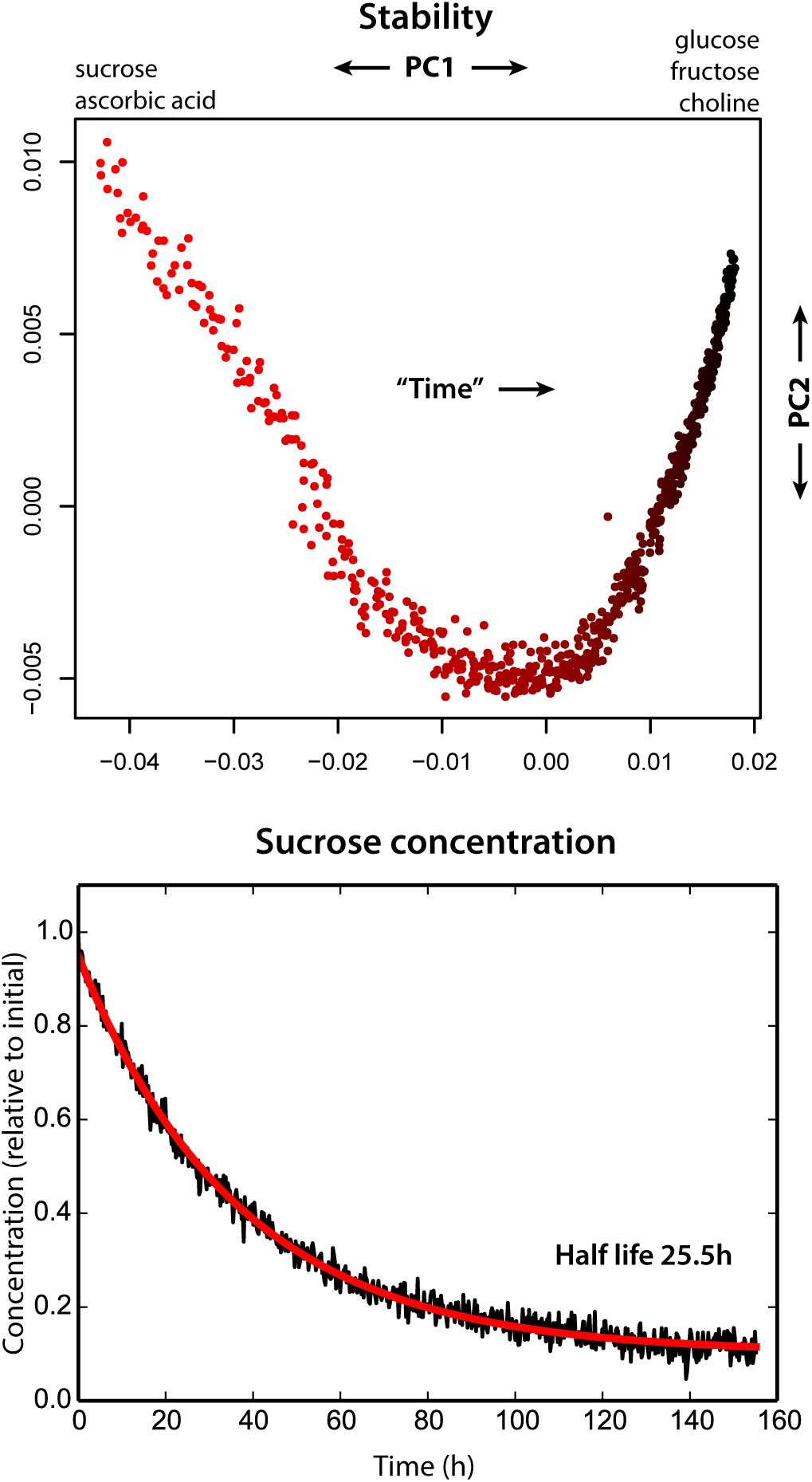
Stability analysis of RIL sample 472 using PCA. Time is represented by color, going from light red to black in 874 second intervals. The “time” marked in the plot is not an actual axis, but the PC1 seems to separate the samples very well according to elapsed time.

## 4. Conclusions

ImatraNMR and SimpeleNMR are free tools for quantitative NMR analysis, which can be used for efficient processing of quantitative NMR spectra. Using these tools, analyzes of both ^1^H and JRES spectra acquired from different lines of *Arabidopsis thaliana* extracts were demonstrated. The results indicate that a combination of JRES spectra, histogram binning (ImatraNMR) and PQ normalization seems to produce suitable data for statistical analysis, yielding superior results compared to basic 1D ^1^H spectra and equidistant binning. This is in line with previous work [69, 58]. JRES spectra seems to work providing more resolution in ^1^H spectra, with the drawback of strong dependence of transverse relaxation rate due to the employed apodization. Relative concentration differences can be observed, but changing field strength, solution viscosity or any other factor affecting T_2_ relaxation rate can alter the measured values.

Several metabolite differences could be identified with multivariate statistical analysis. Especially epithionitriles and iberin nitrile concentration were shown to differ in the main plant lines. In addition, changes during oxidative stress (ozone exposure) were detected, with GABA found as a clear marker for ozone incurred damage. Sample stability was also examined by measuring repetitive ^1^H spectra for extended period of time, with some unstable metabolites found (sucrose, ascorbic acid). These compounds are still sufficiently stable for quantitative analysis, provided that the measurements are done within few hours after sample preparation.

## 5. Acknowledgments

The authors thank Dr. Kirk Marat for correspondence while writing Bruker file handling in SimpeleNMR and Dr. Wolfgang Bermel from Bruker BioSpin for providing insight on implementation details. Dr. Koichiro Shimomura is acknowledged for providing NMR spectrum of C5 epithionitrile. Funding was provided by the Academy of Finland (grants: 135751, 140981, and 273132 to M.B. and the Academy of Finland Center of Excellence in Primary Producers 2014–2019). V.M. and J.H are members of the doctoral programme in chemistry and molecular sciences (CHEMS), while L.V. is a member of the Integrative Life Science Doctoral Program (ILS).

