## Supplementary material for "Automated Processing and Statistical Analysis of NMR spectra obtained from *Arabidopsis thaliana* Extracts"

Valtteri Mäkelä, Lauri Vaahtera, Jussi Helminen

**6 Ozone exposure** **17**

### 1 SimpeleNMR processing and plotting

SimpeleNMR configuration files used to process the different spectra. SimpeleNMR itself was launched with:

```
$ simpele.py d vn fid/1H_ab -o spectra -c simpele/cvi.sconf
```

#### 1.1 Configuration for $^{1}H$

```
zerofill = default
apo_lb = 0.5
phase = auto
autophase_mode = 4

baseline_corr = 1
autophase_blcorr = 1

align = -1 -> 1 @ 0
```

#### 1.2 Configuration for $^{1}H$ JRES

The `project_f2` was removed to produce 2D spectra.

```
proc = 2d

apo_sine = 1.0
apo2d_sine = 1.0

zerofill = default
zerofill2d = default

force_f1freq = y
rotate45 = y
fold = y
project_f2 = sky
```

#### 1.3 Plotting

Plotting example using `simp_view.py` with various options. JRES and 1D $^{1}$H spectra are overlaid, and the `-iil` and `-iil_ind` options are used to give integral area file (from ImatraNMR) and mark some of the areas (found with PCA, associated with epithionitriles).

```
$ simp_view.py 1D_sine1010/V120_jres64.isz \
-iil otsjres1d_hg_histogram_hgl.iil \
-iil_ind 18,38,41,57 \
-crop 1.0:3.5 \
-over ../ots_1h/phcorr1_nuked/nuked_V120_qh1.isz \
-scalearea 1:3.27 \
-pdf plotexample
```

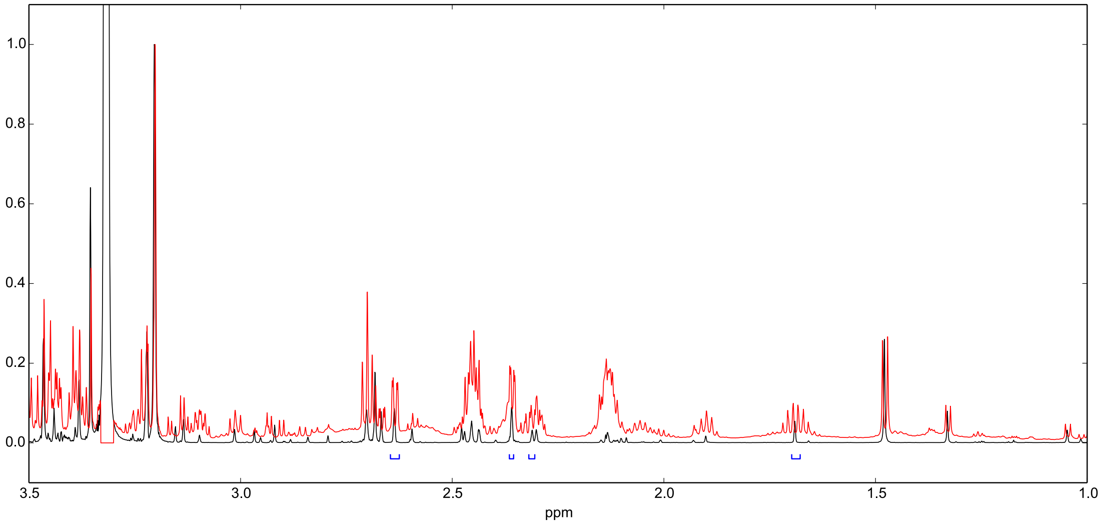

### 2 ImatraNMR processing

ImatraNMR processing scripts, executed with imatra as:

```
$ imatra s imatra/basic_analysis.iscr
```

#### 2.1 Basic $^{1}$H analysis

Sample sets 1A and 1B. Projected 1D JRES analysis was similar.

```
# setup
clear
resetall

var outprefix = basic
var skiptole = 0.0025
var histogtole = 0.0025
var bins = 700

var load_ab = y
var load_cd = y
var export_nuked = n

set outdir out_temp
imgcfg size 1500 400

# load
if $load_ab == y
  load_dir spectra/basic/ab imatra
end
if $load_cd == y
  load_dir spectra/basic/cd imatra
end

# align/scale
align -0.5:0.5 0.0
scale -0.2:0.2

# nuke
nuke 2.19:2.26 # acetone
nuke 3.30:3.33 # MeOH
nuke 4.65:5.12 # water
```

```
# output
set fileprefix ${outprefix}_nuked
set exportprefix nuked

if ${export_nuked} == y
  export is
end

# seek
seekcfg autolimit -1.5:-0.5 20
seekcfg incl 0.5:9
seekcfg skiptole $skiptole
seekcfg histogtole $histogtole
seekcfg
seek

# histogram
seek_histog

# integrate histog
integcfg pqnorm n
integlist_loadhistog
integ

# integrate with pqnorm
set fileprefix ${outprefix}_nuked_pqn
integcfg pqnorm y
integ

# images
imgcfg crop 0.5:9
images seek
images integ

# binning
set fileprefix $outprefix_bin
integlist_clear

# integrate normal
integcfg pqnorm n
integlist_bin 0.5:9.0 $bins
integ

# integrate pqnorm
set fileprefix ${outprefix}_bin_pqn
integcfg pqnorm y
```

```
integ
```

##### 2.2 2D JRES analysis

Sample set 2.

```
clear
resetall

set 2d_mode y

var outprefix = otsjres2d
var htole_d = 0.01
var htole_id = 0.003

imgcfg 2d_limit 0.005
imgcfg 2d_colorf 3
imgcfg crop -0.5:8.0 -0.050:0.050

# settings
set outdir out_temp
set fileprefix ${outprefix}

# load
load2d_dir spectra/ots_jres2d_v2/2D_sine1010_rotsym

# align/scale/blcorr
align -0.5:0.5 0.0 -0.05:0.05 0.0
scale -0.25:0.25 -0.01:0.01
blcorr

# nuke
nuke 3.28:3.34 -0.05:0.05 # MeOH
nuke 4.76:4.85 -0.05:0.05 # water

images

# seek/histogram
set fileprefix ${outprefix}_hg

seekcfg incl 0.5:9.0 -0.050:0.050
seekcfg autolimit -0.9:-0.25 -0.050:0.050 5
seekcfg histogtole ${htole_d} ${htole_id}
```

```
seekcfg
seek

# histogram
seek_histog

# integrate
integcfg pqnorm n
integlist_loadhistog
integ

# pqnorm
set fileprefix ${outprefix}_hg_pqn
integcfg pqnorm y
integ

# images
images integ
images seek
```

#### 3 Model compound syntheses

##### 3.1 Synthesis scheme

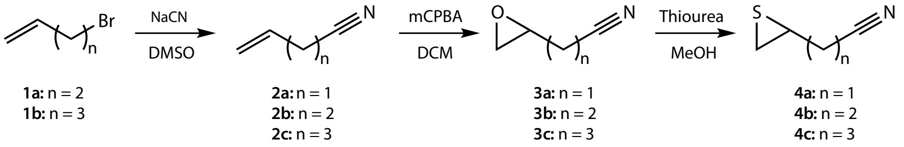

Figure 1: The epithionitrile model compound synthesis scheme.

The epithionitriles **4a** – **4c** were prepared by a modification of a previous method [1]. The starting materials 4-bromo-1-butene (**1a**), 4-bromo-1-pentene (**1b**) and allyl cyanide (**2a**) were obtained from ABCR GmbH, Germany.

##### 3.2 Syntheses

###### 3.2.1 4-Pentenenitrile (2b)

4-Bromo-1-butene (**1a**, 1.00 g, 7.41 mmol) was stirred with dry powdered NaCN (0.400 g, 8.15 mmol, 1.1 eq.) in 17 ml dimethylsulfoxide for 24 hours. The reaction mixture was poured into water (50 ml) and extracted with diethyl ether three times (50 ml total). The ethereal layer was washed with water twice (20 ml total), dried over anhydrous $MgSO_4$ and evaporated to give the title compound as an oil (543 mg, 82% yield of theoretical) in sufficient purity for use without further purification. $^1$H NMR (600 MHz, $CDCl_3$) $\delta$ (ppm): 2.38–2.45 (m, 4H), 5.16 (dd, J = 10.5, 0.9 Hz, 1H), 5.18 (dd, J = 17.3, 0.9 Hz, 1H), 5.83 (ddt, J = 17.3, 10.5, 6.4 Hz, 1H).

###### 3.2.2 5-Hexenenitrile (2c)

The title compound was obtained analogous to 4-pentenenitrile, using 5-bromo-1-pentene (**1b**,1.00 g, 6.71 mmol), NaCN (0.362 g, 7.38 mmol, 1.1 eq.) and 15 ml dimethylsulfoxide, as an oil (594 mg, 85% yield of theoretical) in sufficient purity for use without further purification. $^1$H NMR (600 MHz, $CDCl_3$) $\delta$ (ppm): 1.77 (quin, J = 7.2, 2H), 2.22 (q, J = 7.1 Hz, 1H), 2.35 (t, J = 7.2 Hz, 1H), 5.06 (d, J = 10.1, 1H), 5.10 (dd, J = 17.0, 1.1 Hz, 1H), 5.75 (ddt, J = 17.0, 10.1, 6.7 Hz, 1H).

###### 3.2.3 4,5-Epoxypentanenitrile (3b)

Meta-chloroperbenzoic acid (2.59 g, 67% pure, 10 mmol, 1.5 eq.) was dried under vacuum and dichloromethane (12 ml) and 4-pentenenitrile (**2b**, 543 mg, 6.7 mmol) were added to the flask. The mixture was stirred and heated in an oil bath at 45°C for 12 hours and then cooled down in an ice bath. A solution of $Na_2SO_3$ (830 mg) and $Na_2CO_3$ (1.47 g) in water (15 ml), and active carbon (120 mg) were added and the mixture was stirred in ice bath for 30 minutes, then 30 minutes in room temperature, filtered through celite and the celite washed with dichloromethane (20 ml). The organic phase of the filtrate was separated and the aqueous phase extracted twice with dichloromethane (20 ml total). The combined organic phases were dried over $Na_2SO_4$ and evaporated to give the title compound as an oil (455 mg, yield 70% of theory) in sufficient purity for use without further purification. $^1$H NMR (600 MHz, $CDCl_3$) $\delta$ (ppm): 1.80 (sex, J = 6.9 Hz, 1H), 2.02–2.08 (m, 1H), 2.50 (dd, J = 7.5, 7.0 Hz, 1H), 2.60 (dd, J = 4.6, 2.6 Hz, 1H), 2.85 (dd, J = 4.6, 4.1 Hz, 1H), 3.04–3.07 (m, 1H).

##### 3.2.4 3,4-Epoxybutanenitrile (3a)

The title compound was obtained analogous to 4,5-epoxypentanenitrile, using allyl cyanide (**2a**, 2.00 g, 29.8 mmol), meta-chloroperbenzoic acid (11.52 g, 67% pure, 44.7 mmol, 1.5 eq.) in dichloromethane (50 ml) for 44 hours, as an oil (1.42 g, yield 38% of theoretical) in sufficient purity for use without further purification. $^1$H NMR (600 MHz, $CDCl_3$) $\delta$ (ppm): 2.74–2.76 (m, 4H), 2.89 (t, J = 4.2 Hz, 2H), 3.22 (m, 1H).

##### 3.2.5 5,6-Epoxyhexanenitrile (3c)

The title compound was obtained analogous to 4,5-epoxypentanenitrile, using 5-hexenenitrile (**2c**, 0.59 g, 6.2 mmol), meta-chloroperbenzoic acid (2.37 g, 67% pure, 9.3 mmol, 1.5 eq.) in dichloromethane (12 ml) for 2 hours, as an oil (567 mg, yield 82% of theoretical) in sufficient purity for use without further purification. $^1$H NMR (600 MHz, $CDCl_3$) $\delta$ (ppm): 1.52–1.59 (m, 1H), 1.81–1.90 (m, 3H), 2.39–2.49 (m, 2H), 2.51 (dd, J = 4.9, 2.7 Hz, 1H), 2.78 (dd, J = 4.8, 4.1 Hz, 1H), 2.92–2.95 (m, 1H).

##### 3.2.6 3,4-Epithiobutanenitrile (4a)

Due to the facile polymerisation of the title compound the NMR sample was prepared by dissolving 3,4-epoxybutanenitrile (**3a**, 1.9 mg) and thiourea (2.6 mg) in d4-methanol (0.15 ml) and allowing the reaction to proceed at 35 °C for 1.5 hours, then diluting the mixture with $D_2O$ (0.6 ml). Assignments presented in main article.

##### 3.2.7 4,5-Epithiopentanenitrile (4b)

4,5-Epoxypentanenitrile (3b, 175 mg, 1.8 mmol) was added to a solution of thiourea (206 mg, 2.7 mmol, 1.5 eq.) in methanol (12 ml) and stirred at room temperature for 19 hours. The mixture was diluted with water (50 ml) and extracted with diethyl ether three times (total 15 ml) and the combined ethereal extracts were dried over $MgSO_4$ and stored in freezer. Yield was not determined but the reaction proceeded to completion and no side products were observed. NMR samples were prepared by evaporating an aliquot of the ethereal solution and dissolving the residue in NMR solvent. Assignments presented in main article.

##### 3.2.8 5,6-Epithiohexanenitrile (4c)

The title compound was prepared analogous to 4,5-epithiopentanenitrile from 5,6-epoxypentanenitrile (**3c**, 200 mg, 1.8 mmol), thiourea (206 mg, 2.7 mmol, 1.5 eq.) in methanol (12 ml). Assignments presented in main article.

#### 3.3 Spectra

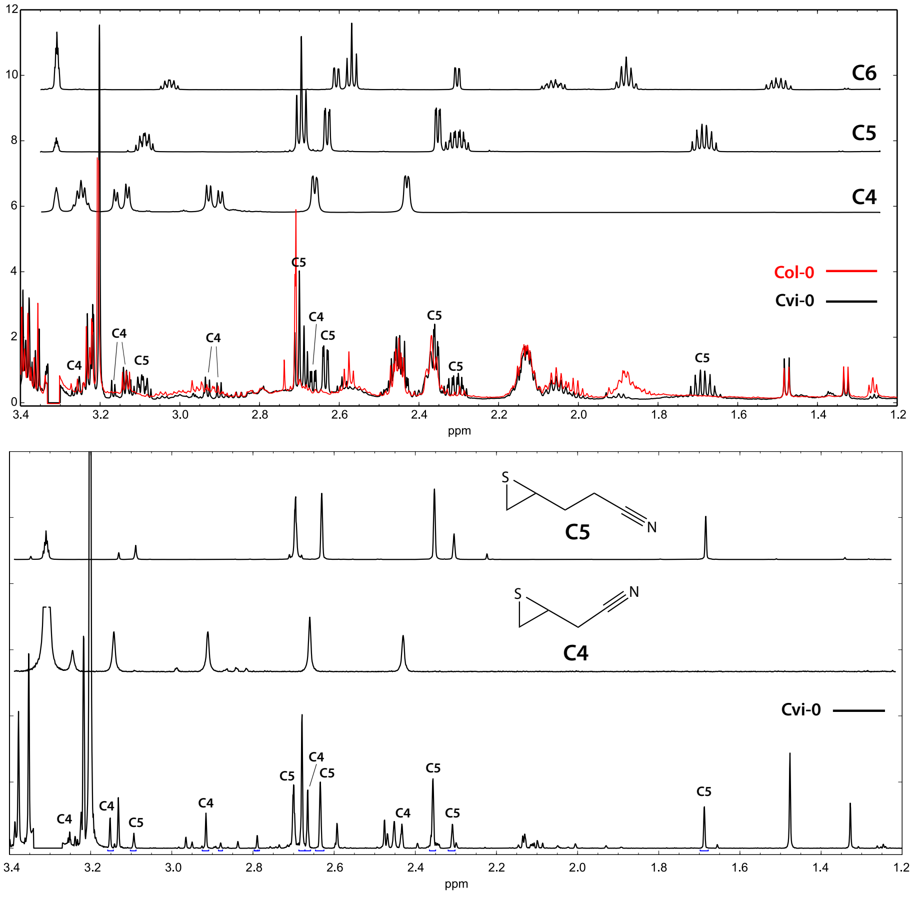

Figure 2: The regular $^{1}H$ (top) and $^{1}H$ JRES (bottom) spectra of different epithionitrile model compounds compared with spectra obtained from Col-0 and Cvi-0.

### 4 RIL analysis

To get an idea on the genetic basis for the differences in nitrile profile, six Cvi-0*Col-0 recombinant inbred lines (RILs) were analyzed. The genome of each RIL is a unique mosaic of the parental genomes, allowing assessment of heritability of a trait. Each analyzed plant line accumulated either C5 epithionitrile or iberin nitrile, but never both. ESM1 and ESP are genes known to be important determinants for plant's nitrile profile, but their origin does not appear to explain the observed differences. The low number of analyzed RILs does not allow the mapping of causative gene(s), but the qualitative nature of this difference implies that there could well be a single causative gene. This preliminary genetic analysis revealed one genetic marker that correlates with the nitrile profile. This marker is located in the south end of chromosome 2, at 18.75 Mb, limiting the potential mapping area to 17.60-19.7 Mb. Further studies would be needed for verification and fine-mapping.

The used RIL lines are described in previous literature [2].

| Sample | ETN integral | ETN verdict | IBN verdict | ESM1 from | ESP from |
|---|---|---|---|---|---|
| 019 | 0.049 | Yes | No | Col-0 | Cvi-0 |
| 359 | 0.076 | Yes | No | Col-0 | Cvi-0 |
| 419 | 0.014 | Yes | No | Cvi-0 | Cvi-0 |
| 424 | 0.041 | Yes | No | Cvi-0 | Cvi-0 |
| 472 | 0.003 | No | Yes | Cvi-0 | Cvi-0 |
| 479 | 0.003 | No | Yes | Col-0 | Cvi-0 |

Table 1: The presence of C5 epithionitrile (ETN) and iberin nitrile (IBN) in the RIL samples.

### 5 Multivariate analyses comparison with alternate processing

Comparing the results of multivariate analysis using equidistant/signal histogram (ImatraNMR) binning and internal standard / PQ normalization.

#### 5.1 PCA

Example command to produce plot:

```
$ simp_mvar.py pca otsjres1d_hg_pqn_integ_sine1010_hgl.csv -pdf pca_hg_pqn \
-group otsjres_groups.txt -noshow -plot_size 120:90
```

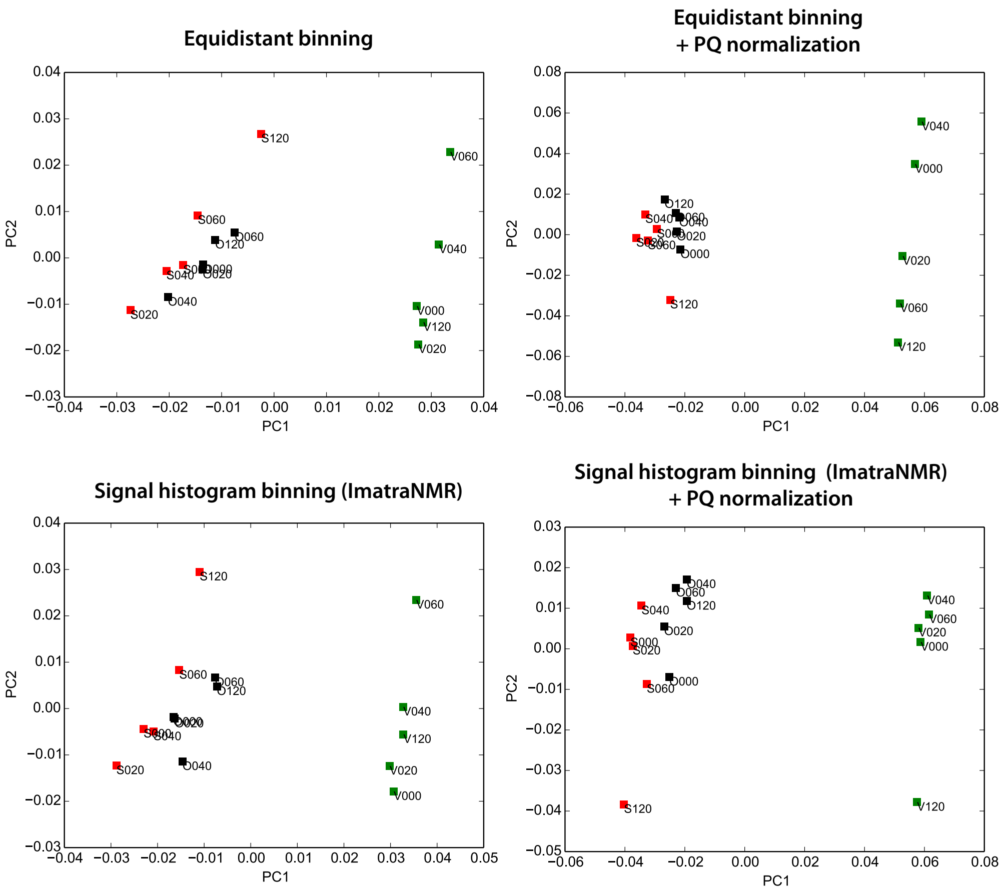

Figure 3: PCA with binning and normalization compared.

#### 5.2 PLS

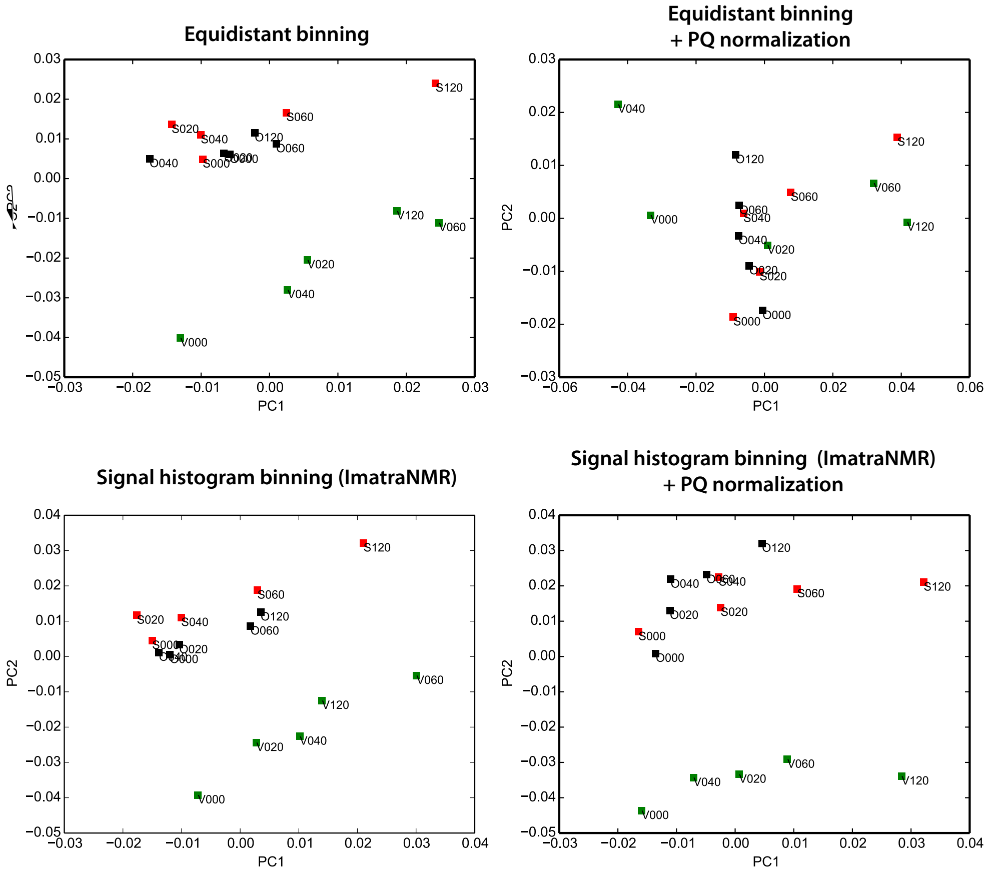

Figure 4: PLS with binning and normalization compared.

#### 5.3 PLS-DA

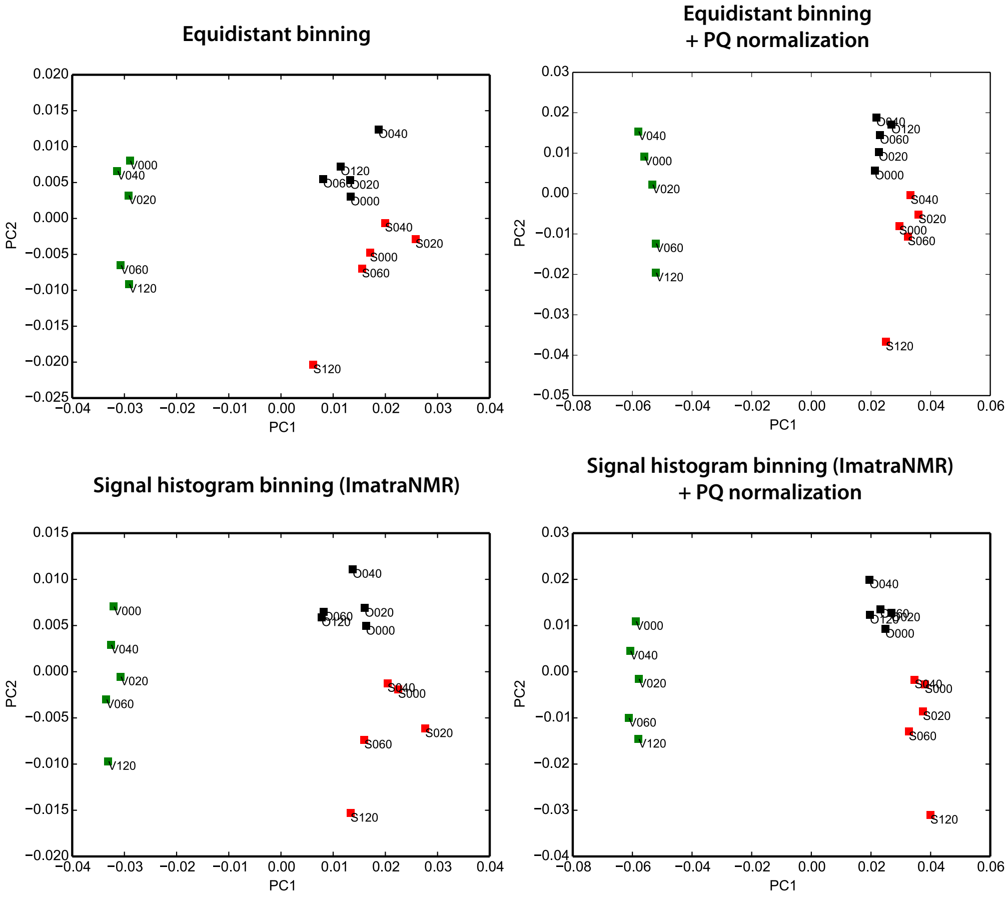

Figure 5: PLS-DA with binning and normalization compared.

### 6 Ozone exposure

Metabolite plots from ozone exposure samples.

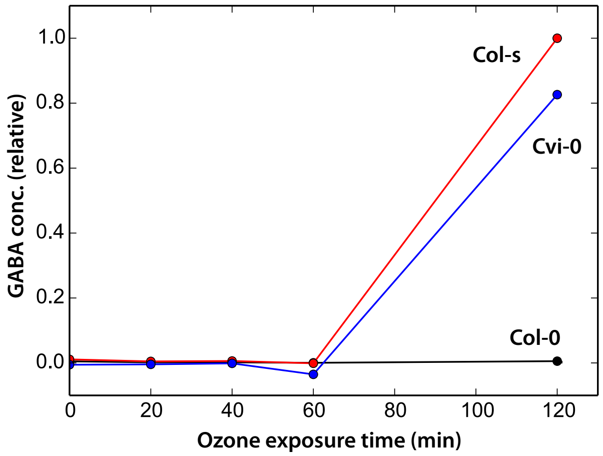

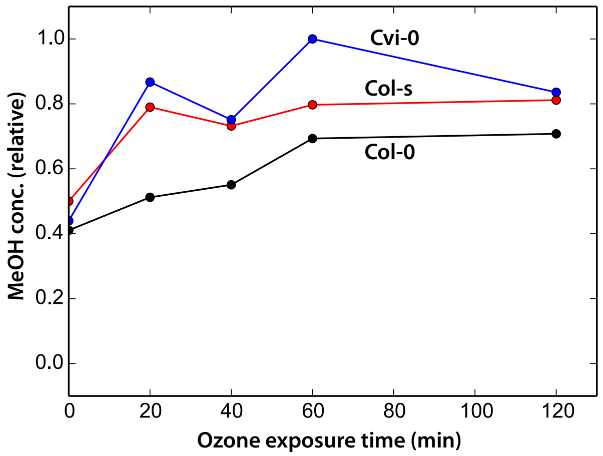
